# Discovering genotype-phenotype relationships with machine learning and the Visual Physiology Opsin Database (VPOD)

**DOI:** 10.1101/2024.02.12.579993

**Authors:** Seth A. Frazer, Mahdi Baghbanzadeh, Ali Rahnavard, Keith A. Crandall, Todd H. Oakley

## Abstract

**Background:** Predicting phenotypes from genetic variation is foundational for fields as diverse as bioengineering and global change biology, highlighting the importance of efficient methods to predict gene functions. Linking genetic changes to phenotypic changes has been a goal of decades of experimental work, especially for some model gene families including light-sensitive opsin proteins. Opsins can be expressed in vitro to measure light absorption parameters, including λmax - the wavelength of maximum absorbance - which strongly affects organismal phenotypes like color vision. Despite extensive research on opsins, the data remain dispersed, uncompiled, and often challenging to access, thereby precluding systematic and comprehensive analyses of the intricate relationships between genotype and phenotype.

**Results:** Here, we report a newly compiled database of all heterologously expressed opsin genes with λ_max_ phenotypes called the Visual Physiology Opsin Database (*VPOD*). *VPOD_1.0* contains 864 unique opsin genotypes and corresponding λ_max_ phenotypes collected across all animals from 73 separate publications. We use *VPOD* data and *deepBreaks* to show regression-based machine learning (ML) models often reliably predict λ_max_, account for non-additive effects of mutations on function, and identify functionally critical amino acid sites.

**Conclusion:** The ability to reliably predict functions from gene sequences alone using ML will allow robust exploration of molecular-evolutionary patterns governing phenotype, will inform functional and evolutionary connections to an organism’s ecological niche, and may be used more broadly for *de-novo* protein design. Together, our database, phenotype predictions, and model comparisons lay the groundwork for future research applicable to families of genes with quantifiable and comparable phenotypes.

**Key Points:**

- We introduce the Visual Physiology Opsin Database (VPOD_1.0), which includes 864 unique animal opsin genotypes and corresponding λ_max_ phenotypes from 73 separate publications.
- We demonstrate that regression-based ML models can reliably predict λmax from gene sequence alone, predict non-additive effects of mutations on function, and identify functionally critical amino acid sites.
- We provide an approach that lays the groundwork for future robust exploration of molecular-evolutionary patterns governing phenotype, with potential broader applications to any family of genes with quantifiable and comparable phenotypes.

## Introduction

Although critical to progress in drug and vaccine design [1–3], responses to climate change [4–8], and bioengineering [4,9–11], accurately predicting gene function from sequences remains a significant challenge. While there are many ways to elucidate genotype-phenotype relationships experimentally, including deep mutational scanning, and in-vitro heterologous expression with phenotyping, these techniques are often tedious and cost-prohibitive, especially when applied to broad comparative studies of gene families. In addition, accurately predicting the phenotype of a protein using computational methods alone is challenging because of data gaps and the sheer complexity of possible relationships between genes and phenotypes, including epistasis and the non-additive effects of different mutations. Machine learning (ML) is gaining traction for its potential broad biological applications, accessibility, and faster speeds, especially in biological contexts where phenotype data are abundant and quantifiable. Here, classical regression and classification algorithms are sometimes used to train models for phenotype predictions using genotype-phenotype data [12,13], while deep learning models can be used to integrate heterogeneous multi-layered omics and environmental data for establishing higher dimensional genotype-phenotype connections [14,15] or *de-novo* protein design [16]. In broader biological contexts, ML models often inform laboratory experiments to predict directional evolution of diseases and their variants [17–19] or to automate image sorting and animal identification from camera trap data [20–22]. In all cases, ML models can be iteratively improved as data gradually accumulates, making them a worthwhile long-term investment.

Decades of laboratory work have led to significant progress in understanding the genetic basis of phenotypic changes for model gene families such as opsins. Opsins are a family of G-protein Coupled Receptors (GPCR) which bind to a chromophore and absorb photons. Opsins have crucial roles in many organismal functions, including circadian rhythms, phototaxis, and image-forming color vision. A critical opsin phenotype is spectral sensitivity - the range of wavelengths to which a gene or organism is sensitive. The main parameter of opsin spectral sensitivity is λ_max_, the wavelength of light (in nm) with maximal absorbance [23]. Common methods of characterizing spectral sensitivities and λ_max_ include organ-level electroretinograms (ERG) [24–26], cell-level microspectrophotometry (MSP) [27–31], and purification of heterologously expressed opsins followed by spectrophotometry. Different opsins are tuned by changes in amino acid sequences to respond to different wavelengths of light, and many previous studies have expressed experimentally mutated opsins and measured spectral sensitivities to establish genotype-phenotype connections [32–36]. Although other factors sometimes affect spectral responsiveness, opsins provide a rare case where an intrinsic molecular function extends rather directly to organismal phenotypes, especially those involving color sensitivity. Despite opsins being a well-studied system with an extensive backlog of published literature, previous authors expressed doubts that sequence data alone can provide reliable computational predictions of λ_max_ phenotypes [37–40]. Furthermore, only the non-animal, microbial, or Type-1 (T1) opsins [41,42] have been cataloged and used to examine genotype-phenotype predictive power of ML models. The extensive data on animal opsin genotype-phenotype associations remains disorganized, decentralized, often in non-computer readable formats in older literature, and under-analyzed computationally.

Here, we report a genotype-phenotype database for animal opsins called the Visual Physiology Opsin Database (*VPOD*). We used standard literature searches to compile all heterologously expressed animal opsin genes with spectral sensitivity measurements. We used this newly compiled and harmonized database to evaluate ML methods for connecting genotypes and phenotypes. We created eleven subsets of the overall database to examine factors that impact the reliability and performance of ML models and compared ML predictions to phylogenetic imputation [43,44]. We also examined whether ML can predict intragenic epistasis, and we predicted amino acid sites particularly important for changing λ_max_. Using our database of 864 unique opsin sequences and corresponding λ_max_ values, we show ML models trained on opsin data accurately predict the λ_max_ of opsins from genetic data alone [highest R^2^ =0.968 with a lowest mean absolute error (MAE) of 6.56 nm], especially when ample and diverse training data are available. ML also predicts some known effects of epistatic mutations on λ_max_. Finally, ML models identify several sites that cause shifts in λ_max_ (e.g., ‘spectral tuning sites’) and sites known to be structurally important, even in the absence of mutant data in training. These results support the use of ML as a reliable and efficient predictor of λ_max_ for previously uncharacterized opsins, as a tool for identifying candidate spectral tuning sites and epistatic interactions, and as a more general method for linking gene sequences and phenotypes.

## Methods

### Compiling a genotype-phenotype database for animal opsins

We collected λ_max_ data for opsins using typical literature review/search methods, documented in the ‘*litsearch’* table of the *VPOD* database to record search engine, keywords, and date of access. We cataloged all usable papers with λ_max_ data in the ‘*references’* table of *VPOD*, recording DOI and a key to link to the search that found the paper. We documented the details of heterologous expression experiments in the ‘*heterologous’* table, including species, GenBank accession number for the sequence, mutation(s) (if applicable) using a machine-readable notation, λ_max_, cell type for expression (e.g., HEK293, COS1, etc.), protein purification method, type of spectrum (e.g., dark or difference spectrum), and a key to link to the corresponding literature source. We input opsin genetic data in an *‘opsins’* table, recording opsin gene family names (e.g., long-wave sensitive=LWS, short-wave sensitive=SWS1, etc.). We also included specific ‘*gene names’* (where applicable), phylum, class, species information, accession number, DNA sequence, amino acid sequence, and the database from which sequences were retrieved (e.g., NCBI). We recreated all mutant and chimeric (e.g. one or more transmembrane domains of the mutant copied from a different sequence to replace the original) opsin sequences based on literature descriptions using a pair of Python scripts (*mutagenesis.py* and *chimeras.py*) available on our GitHub (https://github.com/VisualPhysiologyDB/visual-physiology-opsin-db). We added all heterologously expressed opsins from the literature to the current version of *VPOD*, which we call *VPOD_1.0*. We refer to heterologous data as a data subset named *VPOD_het_1.0*, which will allow for future additions to the database to link specific opsin sequences to λ_max_ values established with microspectrophotometry or other methods.

### Training ML models with *deepBreaks*

We performed all data pre-processing, including data extraction, sequence alignments, and formatting, in the Jupyter notebooks ‘*opsin_model_wf_windows.ipynb*’ or ‘*opsin_model_wf_mac.ipynb*, available on GitHub. We used two multiple sequence alignment methods, MAFFT [45] and MUSCLE [46], and a version of both alignments with a Gblocks [47] refinement (for a total of four alignments), all set to their default parameters to begin to test the sensitivity of model performance to different alignments. We then trained various ML models employing a custom version of *deepBreaks* [48], an ML tool designed for exploring genotype-phenotype associations. *deepBreaks* takes aligned genotype data (DNA, RNA, Amino Acid) and some measure(s) of corresponding continuous or categorical phenotype data as input to train ML models. deepBreaks uses one-hot encoding to convert amino acid sequences into numerical values. One consequence of this encoding is any amino acids at a given position in the alignment, which are not present at that position in any training data, will be treated equivalently as unseen. For example, cases of only A and V at a highly conserved site in the training set that are presented with a sequence with T at that site will be considered as no A and no V. The models cannot distinguish the input whether it’s T or other unseen amino acids at that site. The results produced by *deepBreaks* encompass a compilation of 12 regression ML models [48], showcasing ten metrics of cross-validation performance (ranked by R^2^) and a report derived from the top-performing models, which ranks amino acid positions by their relative importance to the model (from 0.0-1.0, with 1.0 being a site with the highest relative importance) for the phenotype in question (λ_max_). We evaluated the performance of algorithms based on their relative ranks to look for patterns in which algorithms performed better for different data subsets and approaches. *deepBreaks* also produces a set of distribution box plots (default is 100) to visualize phenotypes (λ_max_) associated with a particular amino acid identity at a site of interest, ordered alphabetically.

### Understanding model performance using different subsets of the database

We created eleven data subsets with varying levels of taxonomic and gene family inclusivity (Table 1) to test which factors most impact the reliability/performance of ML methods. We used naming conventions that include versioning to improve reproducibility and reliability of individual datasets and models. For example, one subset combines ultraviolet and SWS opsins, which we named *VPOD_uss_het_1.0.* Our convention is to name the subset (in this case USS = ‘Ultraviolet and Short-wave Sensitive’ opsins); name the source of phenotype data (heterologous = het), and record the version number of the dataset (1.0). We also created subsets for medium-and long-wave sensitive opsins (*VPOD_mls_het_1.0)* and all rod (Rh1) and rod-like (Rh2) opsins (*VPOD_rod_het_1.0*). Other subsets use species taxonomy, one for vertebrates (*VPOD_vert_het_1.0)* and another for invertebrates (*VPOD_inv_het_1.0)*. For taxonomic subsets, we considered all sequences from phylum Chordata as ‘vertebrates’ and the rest as ‘invertebrates’. Another subset excludes all mutant opsin sequences, called ‘wild-types’ (*VPOD_wt_het_1.0)*. A final named subset is the whole data set (*VPOD_wds_het_1.0)*.

**Table 1.** Performance Metrics Across Opsin Subsets and Top Performing Models.

| Name | Data Subset Version | # Seqs | Top ML Algorithm | <sup>b</sup>$R^2$ | <sup>a</sup>MAE [nm] | MAPE [%] | <sup>b</sup>MSE | RMSE |
| --- | --- | --- | --- | --- | --- | --- | --- | --- |
| Whole Dataset | *VPOD_wds_het_1.0* | 864 | LGBM | 0.947 | 7.47 | 1.71 | 207 | 13.8 |
| All Wild Types | *VPOD_wt_het_1.0* | 318 | Bayesian Ridge | 0.902 | 10 | 2.18 | 297 | 16.5 |
| All Mutants | *VPOD_mut_het_1.0* | 546 | LGBM | 0.951 | 7.89 | 1.86 | 194 | 13.4 |
| Vertebrates | *VPOD_vert_het_1.0* | 721 | LGBM | 0.968 | 6.56 | 1.49 | 111 | 10.3 |
| WT Vertebrates | *VPOD_wt_vert_het_1.0* | 274 | GBC | 0.961 | 5.46 | 1.18 | 82.1 | 8.36 |
| Invertebrates | *VPOD_inv_het_1.0* | 143 | LGBM | 0.814 | 14.7 | 3.22 | 614 | 23.1 |
| Rods | *VPOD_rod_het_1.0* | 352 | Bayesian Ridge | 0.834 | 3.51 | 0.71 | 27.7 | 5.04 |
| WT Rods | *VPOD_wt_rod_het_1.0* | 157 | GBC | 0.783 | 3.57 | 0.72 | 31.9 | 5.11 |
| MWS/LWS | *VPOD_mls_het_1.0* | 91 | XGB | 0.677 | 8.77 | 1.82 | 317 | 15 |
| UVS/SWS | *VPOD_uss_het_1.0* | 280 | GBC | 0.821 | 8.02 | 2.06 | 200 | 13.6 |
| WT UVS/SWS | *VPOD_wt_uss_het_1.0* | 66 | Adaboost | 0.865 | 7.79 | 1.87 | 152 | 10.6 |
| T1 Opsins | *Karyasuyama_T1_ops* | 884 | Random Forest | 0.804 | 9.41 | 1.76 | 186 | 13.5 |

<sup>a</sup>Mean absolute error (MAE) is in relation to the absolute error $\lambda_{\max}$ predictions and interpreted in the same units of ‘nm’. <sup>b</sup> $R^2$ or mean square error (MSE) are often interpreted as direct measures of comparing/analyzing model performance and used as training loss terms of the objective function - which measures how well the model fits the training data. One has to often balance between this and the regularization term, which controls the complexity of the model. Thus, a high performance is both simple and predictive; a tradeoff referred to as the ‘bias-variance’ tradeoff.

Using various subsets of data, we performed a number of experiments to better understand the performance of ML models in predicting λ_max_. First, to better understand how training data relate to model performance, R^2^, and training data size, we gradually increased the size of training datasets, using the WDS, Vertebrate, WT, and Rod subsets separately, by adding between 15-50 randomly selected sequences at a time, repeating the process three times per data split (Table S1). We then analyzed the fit between the size of training data sets (x-axis) and model performance (y-axis), comparing six non-linear models with AIC to find the model that best explains the observed variation (Figure S2). Second, to understand if ML could predict known phenotypic changes due to experimental mutations, we queried the top performing WT model (which lacks data from artificially mutated sequences) using all experimentally mutated opsins to predict their phenotypes. We plotted these results using *matplotlib* [49] to visualize characteristics of poorly predicted outliers (e.g., taxonomic bias or sensitivity to mutations which caused large shifts in λ_max_ from the WT). Third, we examined the ability of our models to predict λ_max_ of thirty invertebrate opsins not in *VPOD_1.0* because they are only known from physiological studies (Table S3, Figure S4). Here, we collected data both characterized by single-cell microspectrophotometry (MSP) or electroretinogram methods and with expression localized to cell-type by *in-situ-hybridization* (ISH), to link λ_max_ to a specific opsin (the sequences and metadata can be found in ‘*msp_erg_raw.txt’* and ‘*msp_erg_meta.tsv’*, while the resulting predictions can be found under the *‘msp_tests’* folder on our GitHub repository). Finally, we directly compared predictive capabilities of models trained on different data subsets by randomly selecting and removing the same 25 wild-type ultraviolet or short-wave sensitive opsins from the training data of the WDS, Vertebrate, WT, and UVS/SWS models before training and querying the model with those same sequences following training (Table S3, Figure S5).

### Comparing Machine Learning and Phylogenetic Imputation

We compared performance of ML models to phylogenetic imputation, which estimates phenotypes using phylogenetic information [43,44]. Phylogenetic imputation uses maximum likelihood (we will not abbreviate maximum likelihood as ML to avoid confusion with machine learning), assuming Brownian Motion to predict missing phenotypes using a phylogenetic tree, assuming more closely related species or sequences have more similar phenotypes. For the phylogeny, we constructed opsin gene trees in phyML [50], assuming the ‘WAG’ substitution model [51] and a proportion of 0.029 invariable sites, Gamma as a rate across sites model, and four substitution rate classes. We randomly removed 50 opsin sequences, and their corresponding λ_max_ values from each of the training datasets used to train our ML models (with the exception of the smaller MWS/LWS and invertebrate datasets, in which we only removed 15), then estimated the removed λ_max_ values using phylogenetic imputation. We used the phylogenetic imputation sub-module of the *phytools* R package [52] for performing imputation. We compared imputed and actual λ_max_ using regression. Imputation seemed sensitive to input alignment, perhaps caused by very short or zero length branch lengths in the phylogeny, as we could only complete imputation with *phytools* after removing uninformative and heavily gapped regions with Gblocks. To allow direct comparisons of regressions between imputation and ML, we created ML training-data alignments using MAFFT, MUSCLE, and Gblocks in the same way as for imputation and had ML to predict the λ_max_ for the same set of sequences that were used to the phylogenetic imputation method (Table S6).

### Testing ability of ML to account for intragenic epistasis

Functional predictions are often misled by epistasis [37], so we tested the ability of our WDS models to predict the effects of epistatic mutations by randomly selecting three double mutants with previously demonstrated epistatic effects from training data in which double mutants, each single mutant, and wild type sequence are all characterized by heterologous expression. The three epistatic double mutants are all derived from bovine rhodopsins: D83N_A292S, F261Y_A269T, and A164S_A269T. We removed the double mutants from the training dataset but retained single mutants to retain base effects of each single mutation and test whether the model treats the mutations as additive or epistatic. We hypothesized that the many instances of multi-mutant sequences with epistatic effects in the training set would allow the model to account for some level of intragenic epistasis. In a best case scenario, the model predicts both the magnitude and direction of non-additivity correctly. We then ran a separate test where we removed the same double mutants plus their corresponding single mutants so we could observe whether the WDS model would still predict epistatic effects from wild type data alone. We subsequently repeated this same process for the WT and Vertebrate models (Table S7).

### Identifying known spectral tuning sites

In addition to predicting λ_max_, we wanted to identify amino acid sites with strong effects on the phenotype, called spectral tuning sites for opsins. To do so, *deepBreaks* produces an ‘importance report’ of the relative importance of amino acid positions within the sequence relative to the phenotype. This report is generated for each of the top three performing models, with the addition of a column which calculates the ‘mean relative importance’ value of an individual position. We automated the translation of these feature representations of amino acid positions compared to bovine rhodopsin for the sake of interpretability. We also included the amino acid residue identity at each corresponding position, and whether it is in one of the opsin transmembrane domains (TMD). We used this to provide us with a standardized context for analysis of the most significant positions highlighted by the models which we could use to back-reference against the list of characterized mutants and spectral tuning sites in our database. We analyzed the importance report for each model to see what positions it highlighted as most important, with an extra emphasis placed on the output for the WT models since it was the least likely to be biased by the presence of already-known mutant data (Table S8).

## Results

### Data Description: A genotype-phenotype database for animal opsins

*VPOD_1.0* is a new database, available on GitHub and in DataONE that currently includes all heterologously expressed animal opsins. We refer to a subset of the database with only heterologous data as *VPOD_het_1.0*, although for version 1.0, this is synonymous with the entire database. *VPOD_het_1.0* relies on 68 publications, mainly primary sources, with dates ranging from the 1980’s to 2023. The database contains opsin sequences and phenotype data from 166 unique species (counting 35 reconstructed ancestors), including fishes, amphibians, reptiles, mammals, crustaceans, and bivalves. Altogether, *VPOD_het_1.0* contains 864 unique opsin sequences and corresponding λ_max_ values. This includes 318 unique WT opsins and 546 unique experimentally mutated opsins. Of the mutants, 73 are ‘chimeric’, meaning one or more transmembrane domains of the mutant are copied from a different opsin to replace the original. Phylogenetically, *VPOD_het_1.0* is mainly vertebrate opsins (n = 721), with only 143 unique invertebrate opsins. The vertebrate opsins consist of 113 UVS opsins, 167 SWS opsins, 8 MWS opsins, 83 LWS opsins, 237 Rhodopsin (Rh1), and 113 Rhodopsin-like (Rh2) opsins.

Phenotypically, *VPOD_het_1.0* spans a range of λ_max_ values from 350-611 nm. The highest concentration of phenotype values are between 350-375 nm and 475-525 nm (Figure 1), due to the literature bias favoring characterization of UVS/SWS opsins and rhodopsins (Rh1).

**Figure 1.**
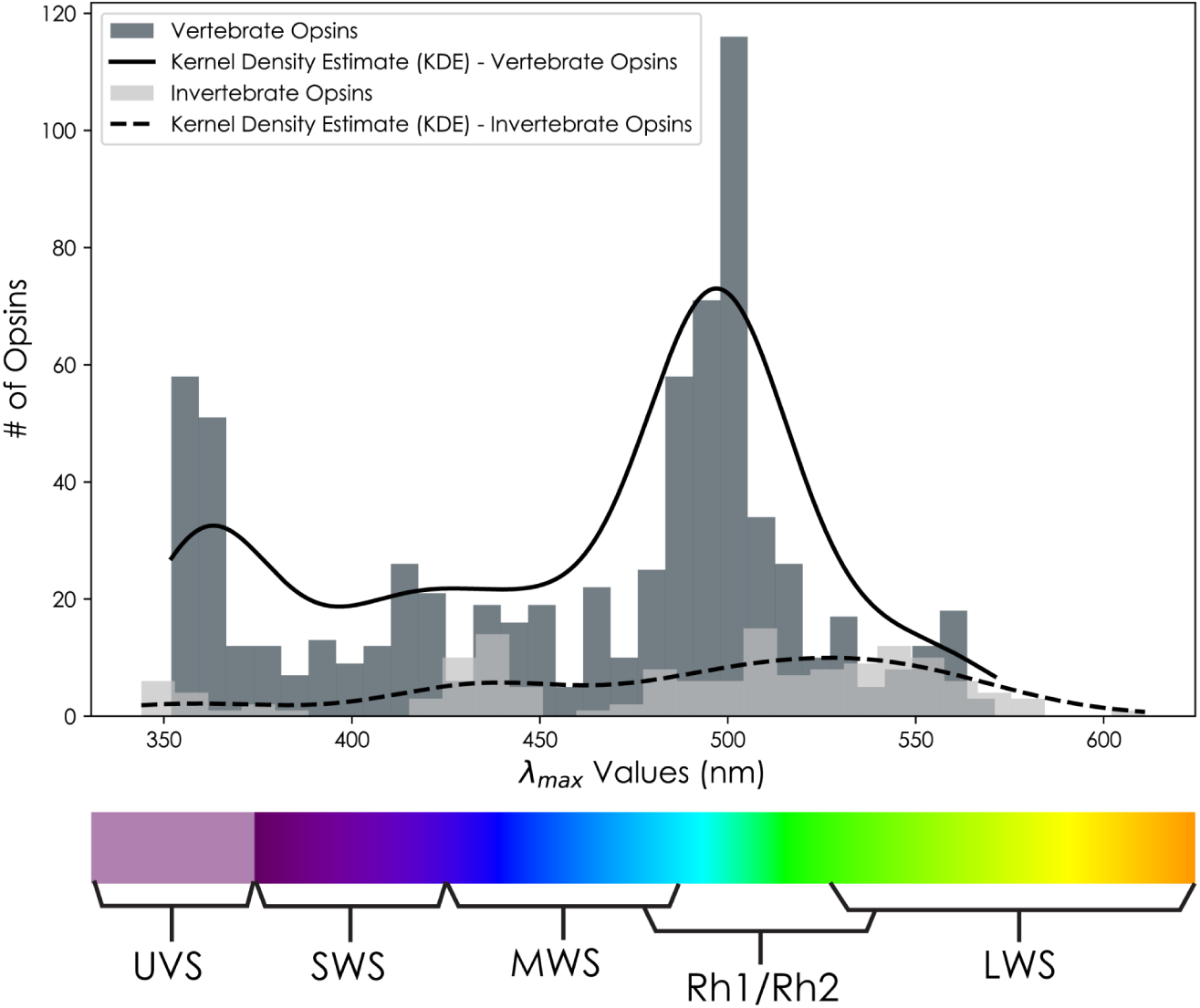
Histogram distributions of Vertebrate and Invertebrate Opsin Light Sensitivity Data - λmax - from *VPOD_het_1.0* with a scaled Kernel Density Estimate (KDE) curves overlaid to better visualize the general shape and characteristics of our λ_max_ distributions. Note an obvious data bias for vertebrate opsins, especially those with λ_max_ values between 350-375nm and 480-510 nm, probably due to focal research on UVS and Rh1 opsins.

### The data used for model training strongly impacts accuracy

Several models trained with different subsets of data predicted λ_max_ with high accuracy (Table 1). The top-performing models from these subsets were also consistently produced using the same five algorithms, including the Gradient Boosting Regressor (GBC)[63,64], Bayesian Ridge (BR)[65,66], Light Gradient Boosting Machine (LGBM)[67], Random Forest (RF)[68], and Extreme Gradient Boosted Machine (XGB)[69]. For example, *VPOD_vert_het_1.0* - trained with all vertebrate wild-type, mutant, and chimeric opsins - had the highest 10-fold-cross-validation (CV) R^2^ (0.968) and lowest mean absolute error (MAE) (6.56 nm) of any models we compared (Figure 2). Similarly, *VPOD_wds_het_1.0*, trained with the whole dataset, had very high R^2^ (0.947) and low MAE (7.47 nm). The two data subsets also shared the same five top performing models (GBC, BR, LGBM, RF, and XGB). In addition, *VPOD_wt_het_1.0* - trained without mutants and only wild type data - had a similarly high R^2^ (0.902) and a low MAE (10.3 nm) when predicting unseen wild type data. Overall, this ‘wild type-only’ model also fared well, even when predicting mutant data not included in the model (Figure 3). At the same time, many instances where mutations caused large shifts in λ_max_ (>10 nm) were not well-predicted by the wildtype-only model, as indicated by large residual values for the predictions of these mutant sequences (Figure 3). Some models, namely those with less than ∼200 training sequences, were far less able to accurately predict λ_max_. For example, *VPOD_mls_het_1.0* – trained only on the 91 MWS/LWS opsins of vertebrates – and *VPOD_inv_het_1.0* – trained only on 144 invertebrate opsins - were among the lowest R^2^ (0.677 and 0.814 respectively) of all models we compared (Table 1). Comparatively, when we trained ML models on the previously published Karyasuyama type 1 opsin dataset we call *Karyasuyama_T1_ops* [41], it produced models with similar performance to the invertebrate model, with a moderate R^2^ (0.804) and MAE (9.41) (Table 1). The similar levels of performances between T1 and invertebrate models were unexpected, especially considering it has a training dataset five times larger than the invertebrate model. One possible explanation is that the very old age of T1 opsins might have led to a higher complexity of genotype-phenotype associations that are not yet well sampled enough to allow good predictions.

**Figure 2.**
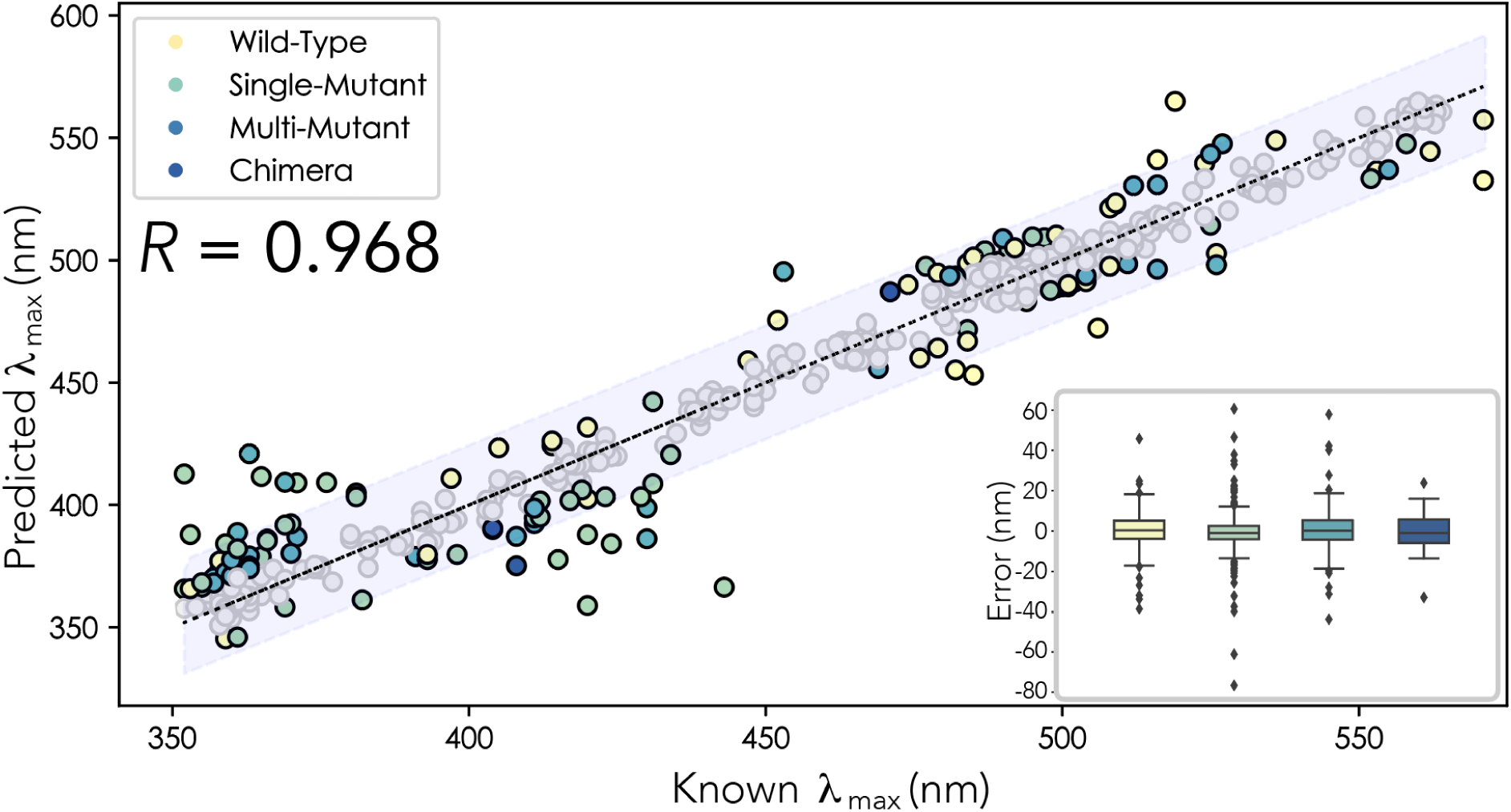
ML model predictions on whole Vertebrate opsin dataset, n = 721, R^2^ = 0.968, MAE = 6.68nm, MAPE = 1.52. Sequences were iteratively and randomly selected to be withheld from the training dataset (n=50) to act as unseen test data. This was repeated until all sequences had been sampled once. Mutant predictions in which the absolute difference between the ‘known’ and ‘predicted’ λ_max_ are <10nm are represented by gray dots. All predictions in which the absolute difference between the ‘known’ and ‘predicted’ λ_max_ are >10nm are represented by colored dots. Yellow dots represent WT predictions, mutants with only a single mutation are green, mutants with greater than one mutation are light-blue, and chimeric opsins are dark-blue. The light-gray bar surrounding the trend-line represents a 95% confidence interval. Inset: Box-plot distribution of prediction error for different opsin data-types from the top performing Vertebrate opsin ML model to better visualize our sources of error. Note, the median for each box-plot hovers around 0nm. Single mutations have the largest spread of error, but this is most likely due to the high abundance of that data-type over all others.

**Figure 3.**
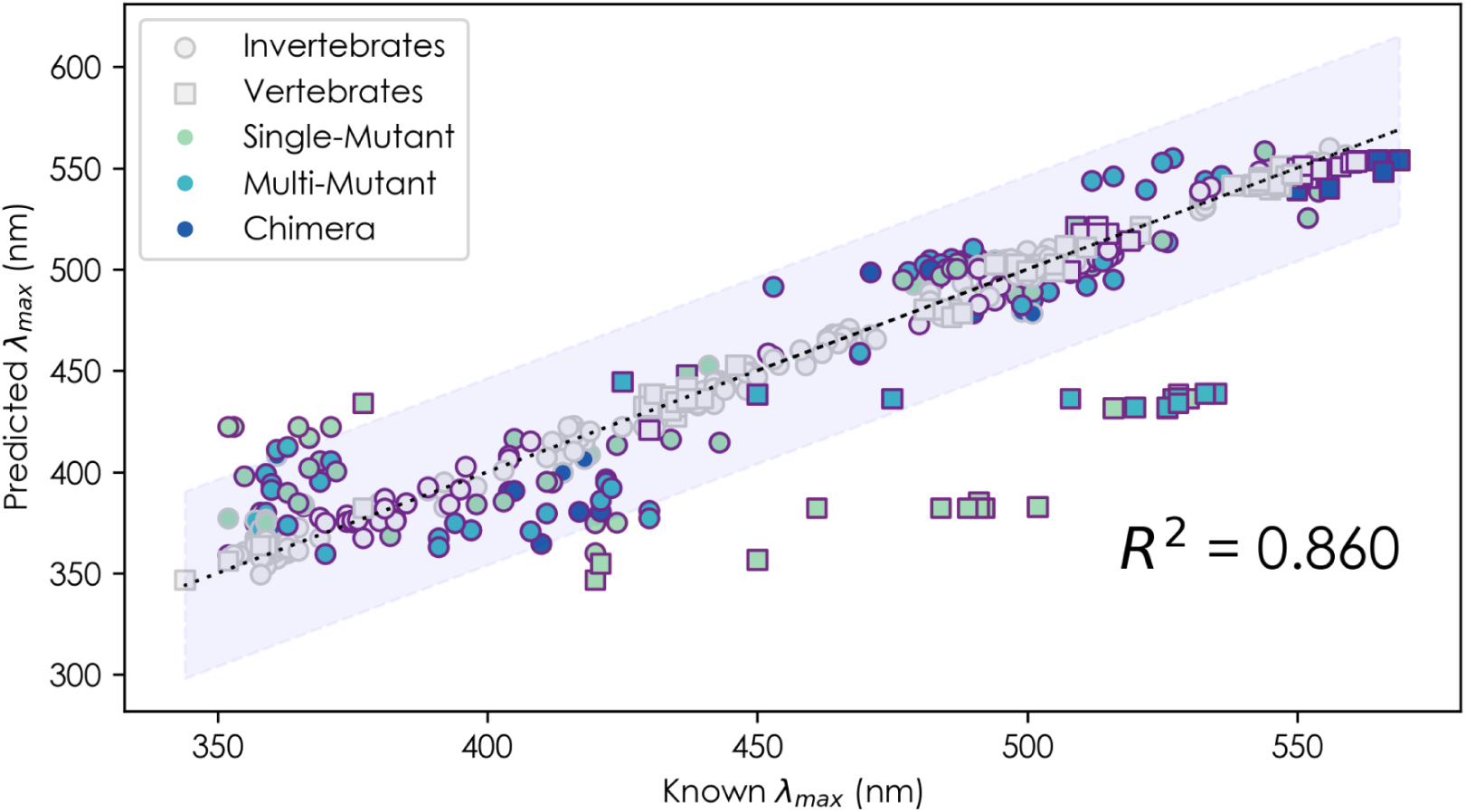
Scatter-plot of Wild-Type Model’s λ_max_ predictions for 547 mutant opsins, with an R^2^ of 0.860, MAE of 12.36 nm, and MAPE of 2.91%. Mutant predictions in which the absolute difference between the ‘known’ and ‘predicted’ λ_max_ are <10nm are represented by gray dots. All predictions in which the absolute difference between the ‘known’ and ‘predicted’ λ_max_ are >10nm are represented by colored symbols, further separated by Invertebrate (Squares) and Vertebrate (Circles) opsins. Mutants with only a single mutation are green, mutants with greater than one mutation are light-blue, and chimeric opsins are dark-blue. Mutations which caused a shift of >10nm from the WT are outlined in purple. The light-gray bar surrounding the trend-line represents a 95% confidence interval.

Data availability improves predictive power, with performance thresholds and plateaus depending on the genetic diversity of the training data. Accuracy in predicting λ_max_ for our models trained on more genotypically and phenotypically diverse subsets of data (WDS, Vertebrate, WT) improves as a function of the number of sequences in a dataset, and shows an initial plateau (R^2^ = ∼0.80-0.90) of diminishing returns around 120-200 sequences, and continues to taper off above 200 sequences (Table S1, Figure S2). For all data subsets, we found the relationship between number of sequences in a dataset and model performance best fits a reciprocal model, which is suitable when the dependent variable plateaus as the independent variable grows larger. We found the coefficients of the reciprocal equations to be different between subsets and to increase in negative magnitude with a decrease in taxonomic/genetic diversity (the Rod model holding the largest negative value of -44). These equations do not account directly for taxonomic, genetic, or phenotypic diversity, as the number of genes is on the x-axis. Therefore, one should be cautious about applying them to predict model performance based on training data size alone.

Although the Rod, UVS/SWS, and MWS/LWS datasets together comprise the training data for the vertebrate model (our highest performing model, R^2^ = 0.968), the individual models trained by these data subsets performed worse than predicted by the trend between number of sequences and model performance observed for the WDS, Vertebrate, and WT subsets (S1, S2). For example, the Rod model, with 352 sequences, should have resulted in a model with an R^2^ around 0.900-0.960 based on the trendlines for the WDS and Vertebrate datasets (S1, S2) but resulted in an R^2^ = 0.831.

When predicting λ_max_ of thirty unseen wild-type invertebrate opsins, almost every model performed rather poorly, with exception of the WT model (n = 30, R^2^ = 0.887, MAE = 17.5) (Table S3, Figure S4). The best performing model produced by the sparsely populated *‘Invertebrate’* dataset could only predict unseen invertebrate opsins with an R^2^ of 0.837 and MAE of 26.3 nm (Table S3. Figure S5). Until the models are trained with more invertebrate (r-opsin) data, we do not put high confidence in the estimates of λ_max_. In contrast, when tested on their ability to predict the λ_max_ for WT SWS opsins, the Vertebrate model (R^2^ = 0.914, MAE = 7.89) outperformed the predictive power of the WDS (R^2^ = 0.833, MAE = 10.2), WT (R^2^ = 0.773, MAE = 9.86) and SWS (R^2^ = 0.788, MAE = 11.6) models, respectively (S3, S5). In other words, the Vertebrate model is both the overall top performing model and the most accurate model for predicting the λ_max_ of SWS opsins.

### ML predictions of λ_max_ are comparable to phylogenetic imputation

Both ML and phylogenetic imputation were often accurate predictors of λ_max_ (Table S6). When using the same test data, ML models usually outperformed phylogenetic imputation, however slightly (S6), albeit using far less computational time, ML using on the order of minutes to calculate models and imputation using on the order of hours to generate opsin phylogenies. The MWS/LWS dataset was the only instance where phylogenetic imputation (R^2^ = 0.784) largely outperformed ML (R^2^ = 0.512). We found our implementation protocol for phylogenetic imputation to require removing aligned sites with extensive gaps using Gblocks, probably lessening impacts on imputation of very short branch lengths. We also used the same alignments for training ML to create direct comparison to imputation results. Interestingly, there was a slight but noticeable decrease in ML performance following Gblocks treatment for the Invertebrate, MWS/LWS, and UVS/SWS datasets (Table S6). The R^2^ of the MWS/LWS model dropped from 0.677 to 0.645, while the Invertebrate model dropped from 0.814 to 0.797 (Table S6). ML performance remained relatively consistent for the WT, Vertebrate, WDS, SWS/UVS, and Rod models, with only a slight reduction in R^2^ (< 0.01) and slight increase in MAE (+/-1nm) for the WT model. The observed disparity in ML performance following Gblocks processing could be attributed to the reduced amount of features in a dataset by removing aligned sites.

### ML often predicts the effects of epistatic mutations

The WDS successfully predicted three out of three instances of epistasis (Table S7) using sequences that were removed from the training data before using the model to predict known epistatic phenotypes. For double mutant D83N_A292S, the model predicted 485.2nm, which was 0.2 nm off the known λ_max_ of 485 nm. If the WDS model believed the sites were additive, the resulting λ_max_ based on adding shifts of single mutants would have been 477.5 nm. Second, for mutant F261Y_A269, the model predicted 520.0nm, for which the known λ_max_ was 520 nm. An additive prediction would have been a λ_max_ 524nm. Third, for mutant A164S_A269T the model predicted a λ_max_ of 515.5 nm, where the known λ_max_ was 514 nm. This is a special case in which the double mutant experiences a form of epistasis where the effect of mutation A269T (λ_max_ = 514) masks the shift otherwise caused by mutation A164S (λ_max_ = 502). Thus, the model seems to have correctly predicted an instance of epistasis in which one mutation masks the effect of another. We believe these results are strong evidence of the model’s eventual capabilities to predict and potentially incorporate the intricacies of intragenic epistasis reliably.

We also queried the WT model with these same double mutants to test the importance of mutant sequences in informing the model on epistatic interactions. However, without any mutant data at all, the WT model did not display the same abilities to predict epistasis in any instance. For the double mutant D83N_A292S, the model predicted neither the individual mutations nor the double mutant would have a significant effect on λ_max_, and all were predicted to be 499.9 nm. For double mutants F261Y_A269 and A164S_A269T, the WT model successfully predicted all individual mutations would cause a red shift, although F261Y and A269 were >3nm off their known λ_max_, but incorrectly treated the mutational effects as additive for the double mutant (Table S7).

Even without information in the training data on the effect of the single experimental mutations of a pair of experimental mutations with known epistatic effects, we could accurately predict epistasis, as long as that sequence variation was present in the training data from wild type sequences. We removed the single mutants F261Y and A269T from the training data. The WDS model predicted a λ_max_ of F261Y as 508.4 nm: 1.6 nm off from the known λ_max_ of 510 nm. The model predicted a λ_max_ of 510.8 nm for mutant A269T: 3.2 nm off its known λ_max_ of 514 nm. In short, the model predicted the correct direction of phenotypic shift for the single mutants, and while the magnitude of the resulting shifts was not precisely predicted, a 2-4 nm difference between the known and predicted λ_max_ value is still less than the MAE of 7.47nm for the WDS model. As for the double mutant F261Y_A269, the model yet again predicted a λ_max_ of 520 nm, even without the direct genotype-phenotype information provided by the single mutants.

### ML predicts tuning sites from Wild-Type sequences alone

The full WT model and its few variants (SWS and Rod WT models) predict several previously characterized ‘spectral tuning sites’ - functionally demonstrated to change λ_max_ - even with no information on mutants used in the training data (Figure 4, Table S8). For the primary WT model alone we found 15 of the top 25 amino acid sites, ranked by relative importance to the model (all ≥ 0.40), were spectral tuning sites previously characterized by mutagenesis and heterologous expression (Table S8). For example, the especially well-characterized position 308 (p308), known for its role in tuning LWS opsins, and considered to be one of the five key sites in characterizing LWS opsins under the ‘Five-Site Rule’ [53], had the highest relative importance value of 1.0 when using the full WT model, indicating the amino acid identity at p308 is especially important for predicting λ_max_. In another example, the full WT model highlighted p181, a phylogenetically conserved counterion in the retinal-opsin Schiff base interaction for all non-vertebrate opsins [54]. Additionally, the transition from E to H at p181 (E181H) is a characteristic of the red-shifted vertebrate LWS opsins [32], easily visualized in Figure 4C. When predicting λ_max_ of bovine rhodopsin with mutation E181H, the WT model predicted a red-shift compared to wild type, as observed with the natural evolution of the LWS opsin lineage. The WT SWS/UVS model similarly highlighted p113, a site functionally characterized as the counterion in the retinal-opsin Schiff base interaction for all vertebrate opsins. Moreover, even the WT Rod model, trained on a mere 157 sequences, identified p292 (S8), another well-characterized and highly conserved spectral tuning site for vertebrate rhodopsins [55–57], as the site with highest relative importance to its predictions of rhodopsin λ_max_.

**Figure 4.**
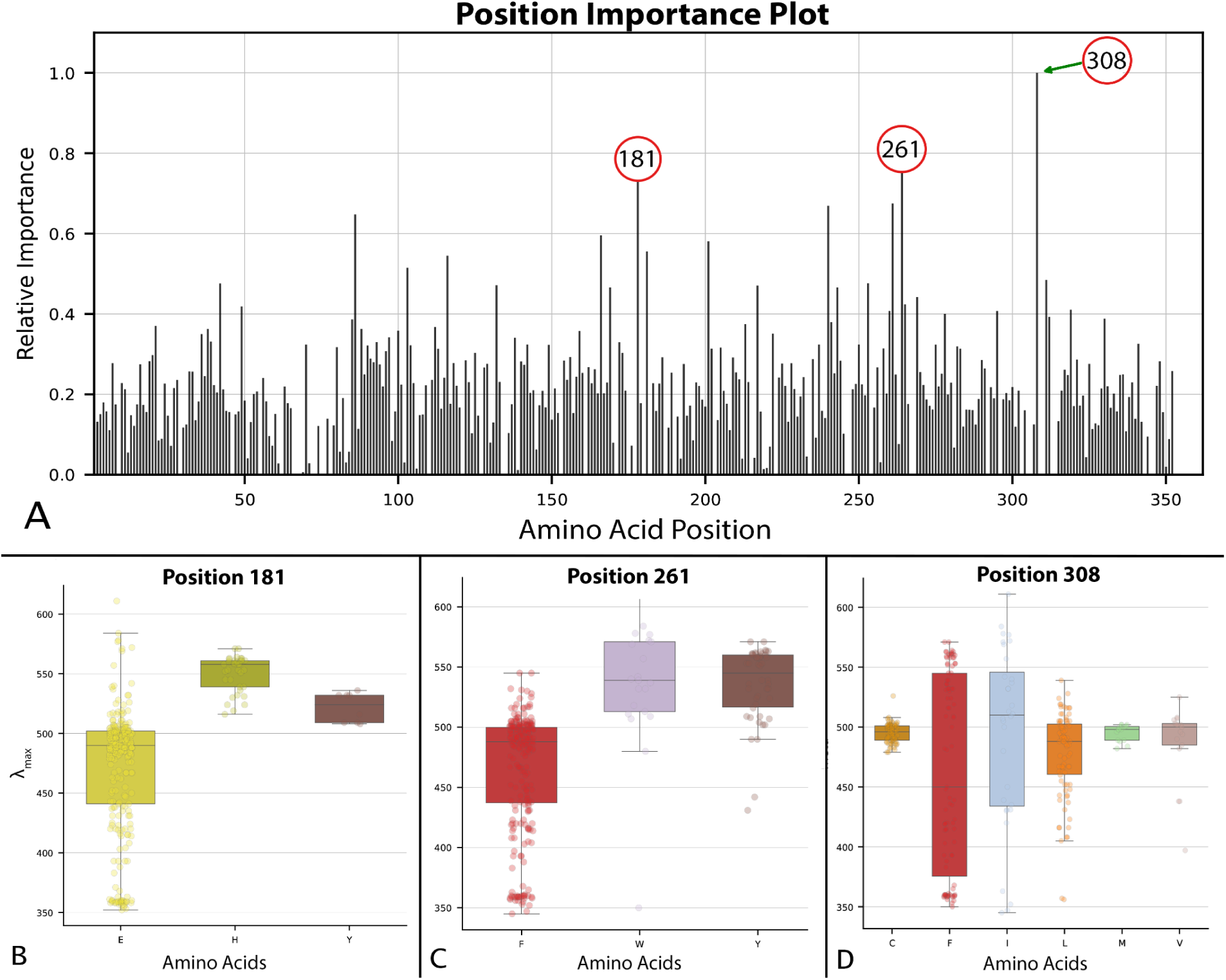
(**A**) Bar graph of relative importance by position generated via ‘BayesianRidge’ ML Regression Model trained on the WT opsin dataset. We interpret positions with higher relative importance as having a larger effect or weight on λ_max_ prediction. Positions 181, 261 and 308 are highlighted because they are among the highest scoring sites and have all been previously characterized as functionally important to opsin phenotype and function. (**B, C, & D**) These distribution box plots provide a visualization for which amino acid (aa) residues at a particular site are associated with different ranges of lambda max at a site of interest, ordered alphabetically, not by frequency (left to right).

## Discussion

To better understand methods to connect genes and their functions, a fundamental goal of biology, we initiate *VPOD,* a database of opsin genes and corresponding spectral sensitivity phenotypes, as a model system for genotype-phenotype interactions. Here, we used *VPOD_1.0* to examine the ability of ML models to predict functions of opsin genes, predict intragenic epistasis, and identify amino acid sites critical for functional changes. In all cases, ML shows promise, especially when given enough training data.

### The important relationship between data availability and predictive power

The predictive power of λ_max_ is generally high when using ML for opsins and it improves with a greater amount and variety of data, albeit with diminishing returns. In particular, the number of opsin genes, their genetic diversity, and the relationship between genetic and phenotypic differences are all critical in determining predictive power. Particularly illustrative of these ideas are our analyses with and without experimentally mutated opsins. Even though we might conceive of all wild type data as natural mutants chosen by evolution, experimentally induced mutations are particularly important by often changing just one amino acid that drastically changes phenotype. As such, we found that including mutant data usually improved predictive power and conversely, predicting some phenotypes from laboratory mutagenesis was sometimes difficult without including other mutant data in model training. Nevertheless, the genotype-phenotype landscape may be sampled well-enough using high numbers of only wild type genes, as evidenced by the small difference in performance when adding mutant data to the wild type subset of well-sampled vertebrate opsins (Table 1). In contrast, adding mutant data to the sparsely sampled invertebrate opsins made a big difference. Here, using only wild type data (ignoring all mutants) led to some very inaccurate predictions, especially of large phenotypic shifts caused by experimental mutagenesis of invertebrate opsins (Figure 3), suggesting the genotype-phenotype space is still undersampled for invertebrates. Therefore, targeting invertebrate (rhabdomeric) opsins should be a high priority for new additions to *VPOD*.

A large diversity of training data is also critical for reliably predicting intragenic epistasis – the non-additive effects on a phenotype of interactions between two or more mutations within a gene – which is common [10,37,39,40,58,59] and an obstacle to connecting genotypes and phenotypes [37,60–62]. Our most complete datasets (whole dataset and vertebrate data) identified known cases of intragenic epistasis, but our models trained without experimental mutagenesis data did not. Similarly to the overall predictive power of λ_max_ above, predicting epistasis probably requires sufficient variation at interacting sites, which seems especially enhanced by experimentally mutated genes.

Variation in the availability of genotype-phenotype data for training not only impacts predictive power of phenotype, but also the converse; the ability to predict amino acid sites that change λ_max_. Several models, including those trained with the whole data set (WDS), Vertebrate, and wild type (WT) data were able to successfully predict previously characterized spectral tuning sites. This is less surprising for models trained with WDS and Vertebrate datasets, due to the prevalence of data, even including mutants in the training data from experiments which specifically targeted sites thought by researchers to be functionally informative. Yet even without any targeted mutational data, three model variants using only wild type data predicted experimentally well-characterized spectral tuning/functional sites, including sites important to the stability of the opsin-chromophore interaction (P181 and P113). This demonstrates the strong potential for ML models to identify amino acid sites that govern phenotype, leading to predictions of candidate spectral tuning sites (CSTS), which can be confirmed with mutagenesis experiments [36,56] if not done so already.

### ML algorithm type contributes to the predictive power of ML models

While probably not as important as the training data used, the ML algorithm itself also impacts predictive power. All five of the best performing ML algorithms (GBC, BR, LGBM, RF, and XGB) are variants of the decision tree model architecture (Table S9), and three out of five, including GBC, LGBM, and XGB, are ‘gradient boosted’ decision tree based ML algorithms. The gradient boosted algorithms all share the same general principles of gradient boosting [63,70] including the use of ensembles of ‘weak learners’, usually decision trees, which work sequentially and ‘gradient descent’ when minimizing a loss function, to improve ML model performance. While LGBM generally performed best for predicting phenotype, it was not as effective in predicting the epistatic effects of mutations, where GBC and XGB showed the highest performance. This suggests that while LGBM excels in general phenotype prediction, the details of GBC and XGB may be better suited for epistasis prediction. The difference likely arises from the unique aspects of each algorithm’s model training and settings of hyperparameters. XGB and LGBM differ from GBC by the addition of a regularization term to the objective function and in the process of ensemble tree construction during model training: GBC and XGB use level-based tree fitting while LGBM uses leaf-based tree-fitting. One consequence of leaf-based tree construction is that due to its faster convergence/training time, it can be more prone to overfitting, as it constructs trees on a ‘best-first basis’ with a fixed number of n-terminal nodes. This creates a model that often performs well but may overgeneralize, missing finer grained collinearities and interdependencies, which would be important for predicting epistasis. As such, our models might be improved by fine-tuning hyperparameters (e.g., learning rate, max-depth, and number of estimators), and the choice of which model to use will depend on the end goals of the analysis.

### The assumptions of our method and limitations of ML extrapolation

Understanding the limitations and assumptions inherent in predictive modeling is vital for accurately interpreting animal color sensitivity from opsin sequences, especially considering the impact of various factors on sensitivity beyond the opsin itself across multiple levels of biological organization. At the cell level, we assume that λ_max_ measured in cell culture (e.g., HEK293, COS cells) is the same as in living photoreceptor cells. We also assume the photopigment uses 11-cis-retinal, as all heterologously expressed opsins in *VPOD* were reconstituted using this chromophore. However, this assumption is violated in some organisms because they use 13-cis-retinal as the *in-vivo* chromophore [71–73], which is associated with a red-shift in λ_max_ [32,71]. At the organ-level, filters such as oil droplets in bird eyes [74–77], pigments in butterfly eyes [78], or a combination of transmissive filter and narrow band reflector in mantis shrimp larval eyes [79], each may selectively influence light reaching photoreceptor cells and therefore animal color sensitivity. Finally, organismal responses to light involve neural processes, so even if an organism possesses the physiological ability to detect certain wavelengths, it still may not have a use for that ability. Similar considerations for all these assumptions will apply when using ML to infer other functions from other genes. In fact, many genes are more susceptible than opsins (but see [80]] showing the pressure of ocean depth may slightly affect λ_max_ phenotypes) to changes in pH, temperature, and other environmental factors [81], such that databases compiling these gene functions should also record these parameters for use in training ML models.

Perhaps the most important caveat of using ML models to accurately predict phenotype or functional sites is that we assume there is a genotype-phenotype association that we can fit to a function and that our models were trained using ample data to capture these associations. Based on the non-linear fit between size of training data set, and model performance, we estimate that including about 200 sequences (and corresponding λ_max_) from a taxonomically and phenotypically diverse range still provides improvements to model performance. Above 200 sequences, there is still improvement, but at a diminishing rate consistent with a reciprocal model (Table S1, Figure S2). When using ML for predicting functionally important sites, the addition of experimental mutants to training data that cause large phenotypic changes could heavily bias which sites are selected as ‘most important’ and potentially mask the importance of other sites. Here again, providing a diverse set of genotype-phenotype data should allow for the discovery of new functional sites, even when including known mutants in the training data with large phenotypic effects. Additionally, not all mutations will have the same effect on different sequences, especially if they are genetically distant; making it important to consider the level of genetic diversity used to train a model when extrapolating to find potentially important functional sites (i.e., if identifying tuning sites for rhodopsins, then using a dataset of only rhodopsins would likely be the best approach, but if data is sparse or if looking for sites that may largely impact spectral tuning across opsin subfamilies, a genetically and phenotypically broad dataset may be better).

## Conclusion

Using opsin sequence data with *deepBreaks*, we were able to train regression-based ML models to reliably predict λ_max_, often accounting for non-additive effects of mutations on function (intragenic-epistasis), and identifying amino acid sites critical for function. We expect future work will improve these already promising results even further through at least two general directions. First, adding more data to *VPOD* will improve results, especially adding invertebrate (rhabdomeric opsins) data, as technical knowledge improves for expressing these genes [35]. In addition, phenotypic data – besides the in-vitro heterologous expression targeted here – is expansive, including λ_max_ measurements from single-cell microspectrophotometry and electroretinograms, but will take considerable effort to link these phenotypes to specific opsin genes. Second, our models can be improved to take advantage of more information. One important addition should be inclusion of physicochemical properties of the amino acids [82], as implemented with success on a small scale of only 26 amino acid positions of microbial opsins to predict red-shifted phenotypes for optogenetics [83]. Additionally, information on protein structure could be particularly important, such as the distance of an amino acid from the binding pocket of the chromophore [84]. While there are only a few solved crystal structures for opsins [85,86] to provide such data, indirect techniques like homology modeling [87] or neural network-base structural prediction [88] might be usable. Other information about opsins could also be predictive, such as which G-protein the opsin signals to, allowing prediction of which amino acids dictate G-protein specificity. Opsin kinetics [e.g. 89], or even the habitat depth at which the animal lives in the ocean, which not only influences light environment but also alters which amino acids are used in opsins [90], could improve predictive power of the ML models.

## Potential Implications

Given the high performance demonstrated in this paper, current models are already robust enough to allow several applications. First, predicting λ_max_ will often be useful, especially for vertebrate opsins. For example, ML could provide an estimate of λ_max_ in a hogfish, whose skin expresses an opsin with unknown absorption and where λ_max_ has implications for a conceptual model of chromatophore expansion [91]. Second, estimates of λ_max_ from opsin sequences formed part of an argument that changes in gene expression, not sequence, adapted Amazon fishes to local light environments [92]. On broader taxonomic scales, predictions of λ_max_ from opsin sequences could expand studies of adaptation, molecular, evolution and constraint in comparison to light environments [93]. Another application could be protein design for optogenetics - the use of genetic light sensors to induce and study expression or response pathways [94–96] - including those associated with embryogenesis [97,98], stress and depression([99–101], or neuronal diseases [102,103]. Finally, our models could be used to simulate molecular evolution under a realistic genotype-phenotype landscape. One shortcoming presently for such simulations is that our models are not trained with non-functional opsins, so even non-functional genes would be predicted to have functional λ_max_ values. A solution could be to add large-scale mutagenesis data to the training set, such as that from deep mutational scanning [104]. As the *VPOD* database expands, there will be many applications for ML, and similar techniques can also be applied to other gene families such as luciferases [16,105,106].

## Availability of Supporting Source Code and Requirements

**Project name**: The Visual Physiology Opsin Database (VPOD)

**Project home page:** https://github.com/VisualPhysiologyDB/visual-physiology-opsin-db

**Operating system(s)**: Windows, MacOS, and Linux

**Programming language:** Python

**Other requirements**: Conda 4.9.2, deepBreaks 1.1.2, GBlocks 0.91b, MAFFT 7.520-1, MUSCLE 3.8.31, mySQL workbench 8.0.36, Python 3.9

## Data Availability

The data set(s) supporting the results and all other code used in this article are available in the ‘*Visual Physiology Opsin Database*’ GitHub repository, [cite unique persistent identifier] and at the ‘*Visual Physiology Opsin Database’* GigaDB repository, [pending acceptance].

### Abbreviations

Adaboost: Adaptive Boosting
AIC: Akaike Information Criterion
COS1: Monkey kidney cell line
CV: Cross-Validation
DNA: Deoxyribonucleic Acid
ERG: Electroretinogram
GBC: Gradient Boosting Classifier
GPCR: G-Protein Coupled Receptors
HEK293: Human embryonic kidney cell line
ISH: In-situ Hybridization
KDE: Kernel Density Estimate
LGBM: Light Gradient Boosting Machine
LWS: Long-Wave Sensitive
MAE: Mean Absolute Error
MAPE: Mean Absolute Percentage Error
ML: Machine Learning
MSE: Mean Squared Error
MSP: Microspectrophotometry
MWS: Medium Wavelength-Sensitive
NCBI: National Center for Biotechnology Information
nm: Nanometers
RMSE: Root Mean Square Error
RNA: Ribonucleic Acid
SWS: Short-Wave Sensitive
T1: Type-1 {Microbial Opsins}
TMD: Transmembrane Domain
USS: Ultraviolet and Short-wave Sensitive
UVS: Ultraviolet-Sensitive
VPOD: Visual Physiology Opsin Database
WAG: Whelan and Goldman substitution model
WDS: Whole Dataset
WT: Wild-Type
XGB: Extreme Gradient Boosting
λ^max^: Lambda Max / Wavelength of light with maximal absorbance

## Authors’ Contributions

THO and KAC conceived the study; SAF performed the analysis; SAF provided online documents and software. SAF and THO drafted the original manuscript. All co-authors discussed the results and edited the final manuscript.

## Acknowledgments

We acknowledge funding from the National Science Foundation DEB-2153773 and IOS-1754770 to THO and DEB-2109688 to KAC and AR. We acknowledge use of computational facilities purchased with funds from the National Science Foundation (CNS-1725797) and administered by the Center for Scientific Computing (CSC). The CSC is supported by the California NanoSystems Institute and the Materials Research Science and Engineering Center (MRSEC; NSF DMR 2308708) at UC Santa Barbara. We acknowledge R. Varney for providing technical support and expertise on the phylogenetic imputation experiments. Thanks to V. Scriven for literature searches and data entry. Thanks to A. Singh and S. Yi for advice. We acknowledge Oakley Lab for comments on an early draft of the manuscript.

