## SupplementalMaterials for "Discovering genotype-phenotype relationships with machine learning and the Visual Physiology Opsin Database (VPOD)"

### Supplementary Material

**Supplementary Material 1 (S1):**

Tracking Model Performance vs. Number of Sequences in Training Data

| Data Subset | # of Sequences | R2 - T1 | R2 - T2 | R2 - T3 | R2 - AVG | RD - R2 |
|---|---|---|---|---|---|---|
| WDS | 864 | 0.947 | 0.947 | 0.947 | 0.947 | 0.96 |
| | 814 | 0.941 | 0.943 | 0.946 | 0.943 | RD - AIC |
| | 764 | 0.937 | 0.945 | 0.943 | 0.942 | -162.00 |
| | 714 | 0.935 | 0.946 | 0.940 | 0.940 | |
| | 664 | 0.929 | 0.944 | 0.938 | 0.937 | |
| | 614 | 0.928 | 0.938 | 0.945 | 0.937 | |
| | 564 | 0.919 | 0.936 | 0.936 | 0.930 | |
| | 514 | 0.912 | 0.922 | 0.944 | 0.926 | |
| | 464 | 0.916 | 0.922 | 0.933 | 0.924 | |
| | 414 | 0.920 | 0.918 | 0.932 | 0.924 | |
| | 364 | 0.924 | 0.903 | 0.923 | 0.917 | |
| | 314 | 0.921 | 0.911 | 0.911 | 0.914 | |
| | 264 | 0.919 | 0.888 | 0.911 | 0.906 | |
| | 214 | 0.921 | 0.893 | 0.900 | 0.905 | |
| | 164 | 0.912 | 0.895 | 0.833 | 0.880 | |
| | 114 | 0.879 | 0.895 | 0.862 | 0.879 | |
| | 64 | 0.765 | 0.799 | 0.713 | 0.759 | |

| Data Subset | # of Sequences | R2 - T1 | R2 - T2 | R2 - T3 | R2 - AVG | RD - R2 |
|---|---|---|---|---|---|---|
| Vertebrate | 721 | 0.968 | 0.968 | 0.968 | 0.968 | 0.99 |
| | 671 | 0.963 | 0.966 | 0.965 | 0.965 | RD - AIC |
| | 621 | 0.960 | 0.969 | 0.965 | 0.965 | -160.00 |
| | 571 | 0.955 | 0.968 | 0.961 | 0.961 | |
| | 521 | 0.949 | 0.966 | 0.965 | 0.960 | |
| | 471 | 0.951 | 0.964 | 0.965 | 0.960 | |
| | 421 | 0.952 | 0.961 | 0.966 | 0.960 | |
| | 371 | 0.951 | 0.956 | 0.962 | 0.956 | |
| | 321 | 0.944 | 0.961 | 0.962 | 0.956 | |
| | 271 | 0.942 | 0.952 | 0.951 | 0.948 | |
| | 221 | 0.938 | 0.928 | 0.949 | 0.939 | |
| | 171 | 0.928 | 0.907 | 0.938 | 0.924 | |
| | 121 | 0.896 | 0.910 | 0.911 | 0.906 | |
| | 71 | 0.925 | 0.816 | 0.868 | 0.870 | |

| Data Subset | # of Sequences | R2 - T1 | R2 - T2 | R2 - T3 | R2 - AVG | RD - R2 |
|---|---|---|---|---|---|---|
| Rod | 352 | 0.834 | 0.834 | 0.834 | 0.834 | 0.93 |
| | 337 | 0.825517 | 0.808289 | 0.802864 | 0.812223 | AIC - R2 |
| | 322 | 0.81498 | 0.835739 | 0.83895 | 0.82989 | -122.00 |
| | 307 | 0.864023 | 0.831771 | 0.850227 | 0.848674 | |
| | 292 | 0.83162 | 0.853479 | 0.804649 | 0.829916 | |
| | 277 | 0.832486 | 0.811542 | 0.827711 | 0.823913 | |
| | 262 | 0.852768 | 0.821254 | 0.839961 | 0.837994 | |
| | 247 | 0.798275 | 0.82422 | 0.818958 | 0.813817 | |
| | 232 | 0.838121 | 0.773139 | 0.789483 | 0.800248 | |
| | 217 | 0.799491 | 0.773947 | 0.768506 | 0.780648 | |
| | 202 | 0.840374 | 0.749616 | 0.759987 | 0.783326 | |
| | 187 | 0.812757 | 0.782659 | 0.800555 | 0.798657 | |
| | 172 | 0.772936 | 0.746982 | 0.777117 | 0.765678 | |
| | 157 | 0.740628 | 0.725929 | 0.726127 | 0.730895 | |
| | 142 | 0.727746 | 0.807972 | 0.668367 | 0.734695 | |
| | 127 | 0.692876 | 0.70442 | 0.765842 | 0.721046 | |
| | 112 | 0.703357 | 0.641533 | 0.74384 | 0.696243 | |
| | 97 | 0.628481 | 0.663658 | 0.763486 | 0.685208 | |
| | 82 | 0.453915 | 0.414098 | 0.603912 | 0.490642 | |
| | 67 | 0.105291 | 0.047832 | 0.556672 | 0.236599 | |
| | 52 | -0.09295 | 0.041141 | 0.340028 | 0.096072 | |
| Data Subset | # of Sequences | R2 - T1 | R2 - T2 | R2 - T3 | R2 - AVG | RD_R2 |
| WT | 318 | 0.902 | 0.902 | 0.902 | 0.902 | 0.84 |
| | 303 | 0.873 | 0.911 | 0.899 | 0.892 | RD_AIC |
| | 288 | 0.891 | 0.876 | 0.873 | 0.884 | -89.00 |
| | 273 | 0.890 | 0.866 | 0.868 | 0.878 | |
| | 258 | 0.881 | 0.859 | 0.879 | 0.870 | |
| | 243 | 0.854 | 0.884 | 0.825 | 0.869 | |
| | 228 | 0.851 | 0.875 | 0.873 | 0.863 | |
| | 213 | 0.826 | 0.775 | 0.863 | 0.800 | |
| | 198 | 0.870 | 0.856 | 0.856 | 0.863 | |
| | 183 | 0.866 | 0.840 | 0.849 | 0.853 | |
| | 168 | 0.856 | 0.830 | 0.776 | 0.843 | |
| | 153 | 0.751 | 0.826 | 0.832 | 0.788 | |
| | 138 | 0.744 | 0.785 | 0.806 | 0.765 | |
| | 123 | 0.747 | 0.855 | 0.808 | 0.801 | |

| | | | | |
|---|---|---|---|---|
| 108 | 0.712 | 0.864 | 0.738 | 0.788 |
| 93 | 0.749 | 0.886 | 0.574 | 0.817 |
| 78 | 0.660 | 0.731 | 0.747 | 0.695 |
| 63 | 0.731 | 0.715 | 0.460 | 0.723 |
| 48 | 0.343 | 0.654 | 0.677 | 0.499 |
| 33 | -0.277 | | | -0.277 |

*T = Test , RD = Reciprocal Decay, AIC =  Akaike Information Criterion*

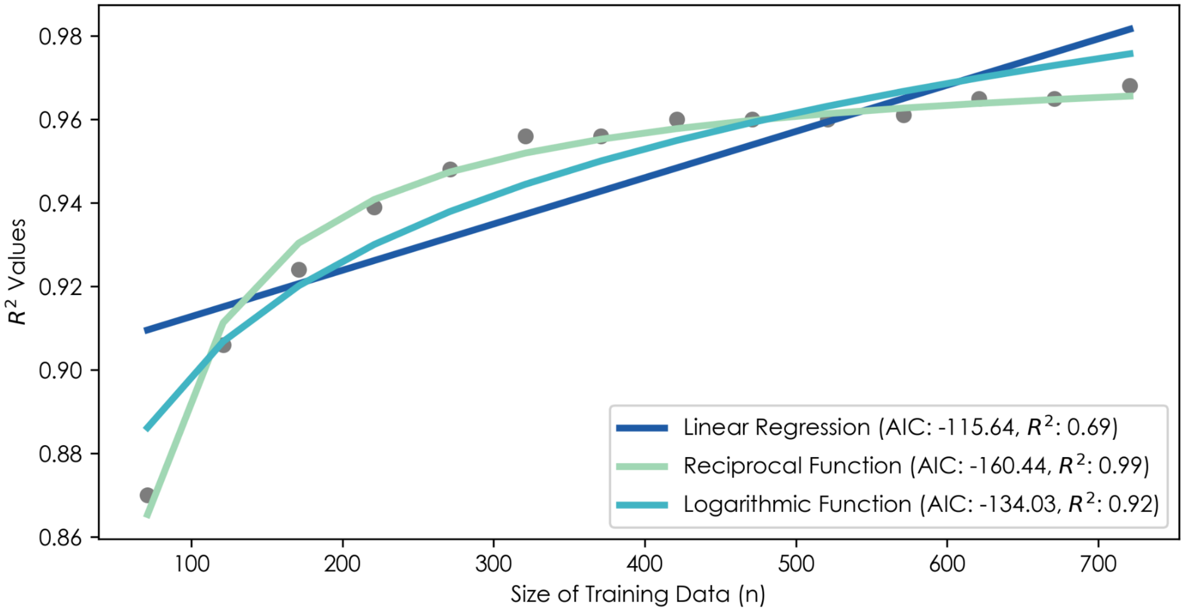


*Supplementary Material 2 (S2): Three functions fitted to visualize the relationship between training data size (number of genotypes and corresponding phenotypes) vs. model performance (R2) based on results from the Vertebrate subset of data. The Akaike Information Criterion (AIC) is a measure used for model selection when comparing different statistical models, accounting for both the goodness of fit of the model and the simplicity of the model (the number of parameters used). The goal is to find a balance between a model's ability to explain the data and its complexity, preventing overfitting.*

**Supplementary Material 3 (S3):**
Comparing ML Predictions on Invertebrate and Vertebrate UVS/SWS Opsin MSP Data

| Data Split | MSP R2 | MAE [nm] | MAPE [%] | SWS Test R2 | MAE [nm] | MAPE [%] |
|---|---|---|---|---|---|---|
| Whole Dataset | 0.704 | 30.6 | 6.8 | 0.833 | 10.2 | 2.45 |
| Vertebrates | 0.044 | 70.8 | 14.6 | 0.914 | 7.89 | 1.9 |
| Invertebrates | 0.837 | 26.3 | 6.95 | - | - | - |
| Wild Types Only | 0.887 | 17.5 | 4.06 | 0.773 | 9.86 | 2.46 |
| UVS/SWS | - | - | - | 0.788 | 11.6 | 2.92 |

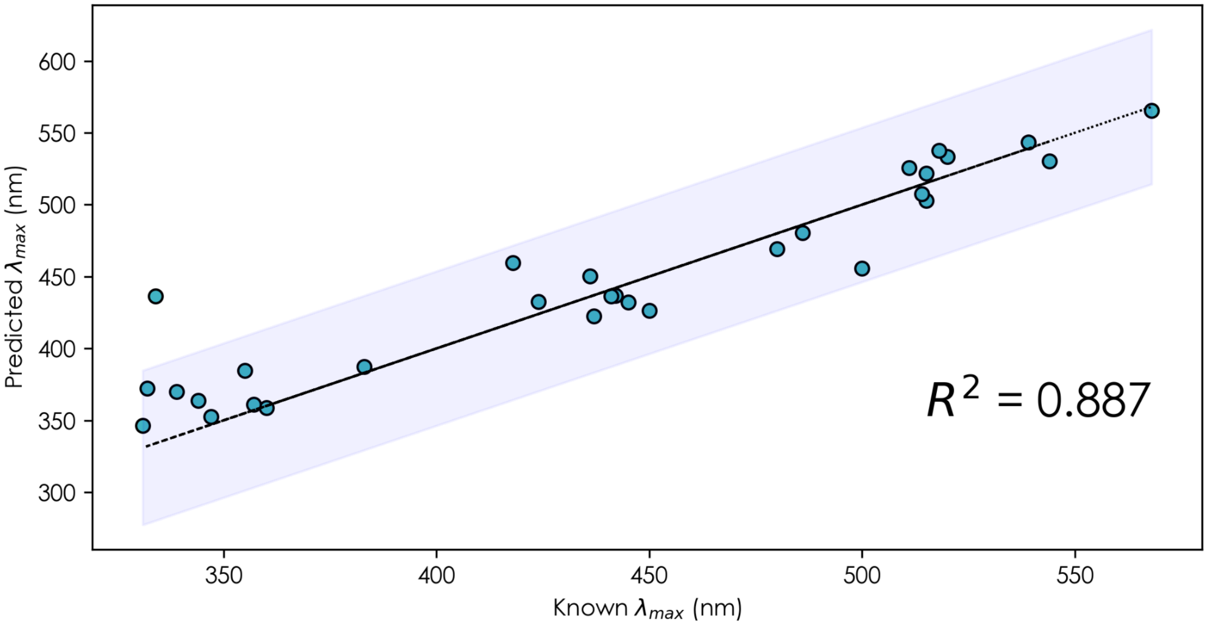


**Supplementary 4**: Graph of WT model predictions for 30 unseen invertebrate opsins, R2 = 0.887, MAE =17.5nm, MAPE = 4.05. All the ‘known’ λmax values are from physiological measures, including MSP or ERG measurements (instead of purified heterologously expressed opsins), and are linked to a particular opsin sequence by in-situ hybridization. The light-gray bar surrounding the trend-line represents a 95% confidence interval.

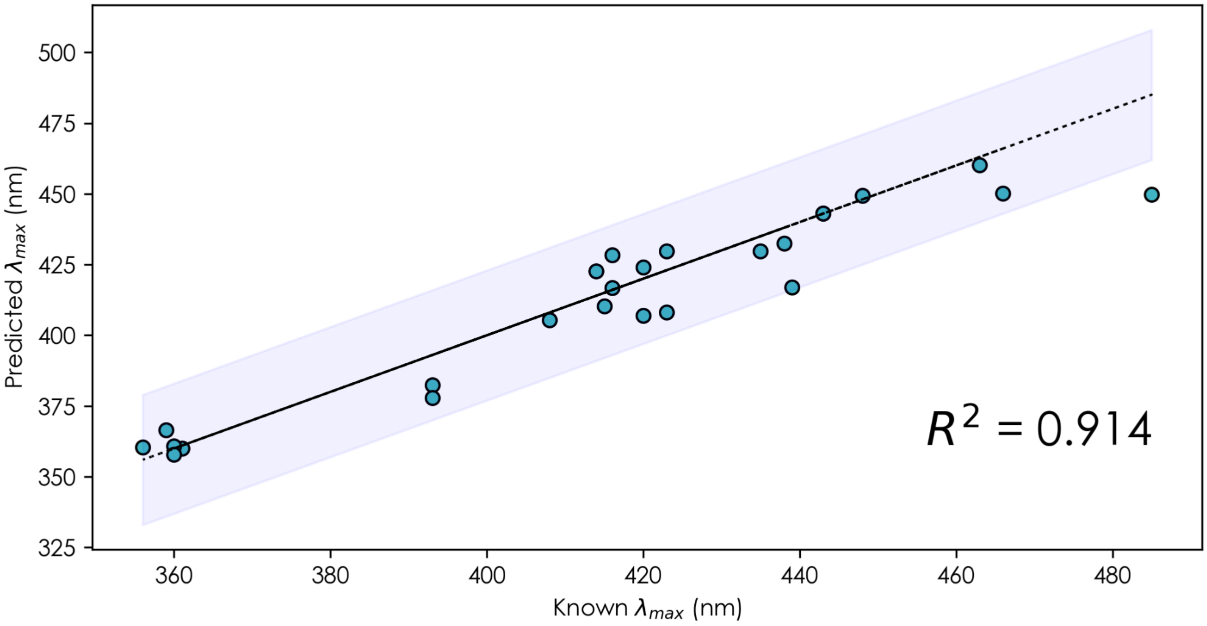


**Supplementary 5 (S5)**: Graph of Vertebrate model predictions for unseen WT-UVS/SWS data, n = 25, R2 = 0.914, MAE =7.89 nm, MAPE = 1.90. All sequences were randomly selected from the UVS/SWS model under the condition that they were WT opsins. The light-gray bar surrounding the trend-line represents a 95% confidence interval.

**Supplementary 6 (S6):**
Comparing Performances of ML Predictions and Phylogenetic Imputation on a Subsample of Opsin Data

| Subset | MAFFT Sample R2 | MUSCLE deepBreaks R2 | MUSCLE Sample R2 | Gblock deepBreaks R2 | Gblock Sample R2 | Imputation Sample Adj. R2 | Imputation . MAE (nm) | Imputation MAPE (%) | Sample Size (n) |
|---|---|---|---|---|---|---|---|---|---|
| Wild-Type | 0.868 | 893 | 0.918 | 0.900 | 0.863 | 0.836 | 8.78 | 1.85 | 50 |
| Vertebrates | 0.968 | 0.967 | 0.972 | 0.967 | 0.967 | 0.949 | 6.76 | 1.48 | 50 |
| WDS | 0.948 | 0.946 | 0.96 | 0.942 | 0.964 | 0.958 | 7.6 | 1.65 | 50 |
| Rods | 0.848 | 0.843 | 0.824 | 0.843 | 0.865 | 0.713 | 4.36 | 0.877 | 50 |
| Invertebrates | 0.706 | 0.814 | 0.711 | 0.797 | 0.678 | 0.758 | 18.3 | 3.43 | 15 |
| UVS/SWS | 0.87 | 0.818 | 0.919 | 0.820 | 0.921 | 0.922 | 10.9 | 2.78 | 25 |
| MWS/LWS | 0.499 | 0.657 | 0.497 | 0.645 | 0.512 | 0.784 | 5.91 | 1.12 | 15 |

**Supplementary 7 (S7) :** Results for epistasis test on the WDS, Vertebrate, WT, and Rod models.

| Mutant | Known λmax (nm) | WDS λmax Prediction (nm) | Vertebrate λmax Prediction (nm) | WT λmax Prediction (nm) |
|---|---|---|---|---|
| NM_001014890 [WT] | 500 | - | - | - |
| NM_001014890_D83N_A292S | 485 | 485.2 | 483.95 | 499.91 |
| NM_001014890.2_F261Y_A269T | 520 | 520.0 | 515.39 | 515.3875 |
| NM_001014890.2_A164S_A269T | 514 | 515.5 | 513.8 | 510.2689344 |

**Supplementary 8 (S8):**
Functionally characterized spectral tuning sites predicted by the WT models

| Data Subset | Position on Bovine | Importance Value (0.0-1.0) | AA Residue on Bovine | TMD | Verified Tuning Site |
|---|---|---|---|---|---|
| All WT Opsins | 308 | 1 | M | 7 | Yes |
| | 261 | 0.675 | F | 6 | Yes |
| | 86 | 0.648 | M | 2 | Yes |
| | 201 | 0.581 | E | CT/EC | Yes |
| | 181 | 0.555 | E | CT/EC | Yes |
| | 116 | 0.545 | F | 3 | Yes |
| | 253 | 0.476 | M | 6 | No |
| | 42 | 0.476 | A | 1 | No |
| | 217 | 0.471 | I | 5 | Yes |
| | 169 | 0.466 | A | 4 | Yes |
| | 243 | | | | No |
| | 49 | 0.419 | M | 1 | Yes |
| | 295 | 0.407 | A | 7 | Yes |
| | 269 | 0.442 | A | 6 | Yes |
| Rod WT Opsins | 292 | 1 | A | 7 | Yes |
| | 122 | 0.417 | E | 3 | Yes |

**Supplementary 8 (S8):**
Functionally characterized spectral tuning sites predicted by the WT models

| Data Subset | Position on Bovine | Importance Value (0.0-1.0) | AA Residue on Bovine | TMD | Verified Tuning Site |
|---|---|---|---|---|---|
| All WT Opsins | 308 | 1 | M | 7 | Yes |
| | 261 | 0.675 | F | 6 | Yes |
| | 86 | 0.648 | M | 2 | Yes |
| | 201 | 0.581 | E | CT/EC | Yes |
| | 181 | 0.555 | E | CT/EC | Yes |
| | 116 | 0.545 | F | 3 | Yes |
| | 253 | 0.476 | M | 6 | No |
| | 42 | 0.476 | A | 1 | No |
| | 217 | 0.471 | I | 5 | Yes |
| | 169 | 0.466 | A | 4 | Yes |
| | 243 | | | | No |
| | 49 | 0.419 | M | 1 | Yes |
| | 295 | 0.407 | A | 7 | Yes |
| | 269 | 0.442 | A | 6 | Yes |
| Rod WT Opsins | 292 | 1 | A | 7 | Yes |
| | 205 | 0.256 | I | 5 | Yes |
| SWS WT Opsins | 113 | 0.302 | T | 3 | Yes |
| | 81 | 0.156 | M | 2 | Yes |
| | 114 | 0.124 | L | 3 | Yes |

**Supplementary 9 (S9):**
Ranked ML Algorithm Performances

| Model Algorithm | Top Model Count | Total Points | Overall Ranking |
|---|---|---|---|
| ***gbc*** | 3 | 120 | 1 |
| ***BayesianRidge*** | 2 | 104 | 2 |
| ***lgbm*** | 4 | 100 | 3/4 |
| ***rf*** | 1 | 100 | 3/4 |
| ***XGB*** | 1 | 87 | 4 |
| ***ET*** | 0 | 68 | 5 |
| ***Adaboost*** | 1 | 51 | 6 |
| ***DT*** | 0 | 50 | 7 |
| ***LassoLars*** | 0 | 48 | 8 |
| ***Lasso*** | 0 | 41 | 9 |
| ***HubR*** | 0 | 27 | 10 |
| ***LR*** | 0 | 0 | 12 |
